# G protein βγ subunits bind to and inhibit the function of multiple Qa- and Qb,c-SNARE isoforms

**DOI:** 10.1101/2022.05.30.494040

**Authors:** Zack Zurawski, Spencer Huynh, Ali Kaya, Karren Hyde, Heidi E. Hamm, Simon Alford

## Abstract

While the ability of G protein βγ subunits (Gβγ) to bind to and functionally inhibit the neuronal SNARE proteins Stx1A, SNAP25, and synaptobrevin in the presence of the calcium sensor synaptotagmin I is well documented, these three SNARE proteins, which form the core SNARE complex for synchronous evoked release in neurons, are but a subset of the larger family of SNARE proteins, which participate in many other exocytic processes within the cell and in other populations of secretory cells throughout the body, from which the release of neurotransmitters, hormones, and other factors is regulated by G_i/o_-coupled GPCRs. The ability of Gβγ to regulate these processes is unknown. To investigate the feasibility of this mechanism to inhibit SNARE function more broadly, we utilized a series of biochemical assays of binding and function with four Qa-SNAREs (Stx1A, Stx2, Stx3, and Stx4) and four Qb,c-SNAREs (SNAP25, SNAP23, SNAP29, and SNAP47) in tandem with the R-SNARE synaptobrevin, synaptotagmin I, and Gβγ. Gβγ was found to bind to multiple Qa-SNARE isoforms as well as SNAP23, and inhibit the lipid mixing function of these SNAREs, as well as SNAP29. Together, this data suggests a more broad role for the Gβγ-SNARE pathway in the regulation of exocytosis beyond cells that express Stx1A or SNAP25.

## Introduction

Inhibitory Gi/o-coupled G-protein coupled receptor signaling occurs via multiple independent pathways involving both Gα_i/o_ and Gβγ. The inhibition of exocytosis by Gi/o-coupled GPCRs has been shown to occur via a well-studied pathway involving the interaction of Gβγ heterodimers with N, P/Q, or R-type voltage-gated calcium channels^1–5^, reducing inward Ca^2+^ flux, with many Gi/o-coupled GPCRs signaling exclusively via this mechanism^6–8^. Gβγ signaling has also been shown to occur downstream of Ca^2+^ entry^9–11^, a process initially identified at the neuronal presynaptic terminal. Gβγ subunits bind to the SNARE complex consisting of syntaxin1A, SNAP25, and synaptobrevin, competing with the calcium sensor synaptotagmin I for binding sites on SNAP25^12–15^. This mechanism has been shown to occur at multiple populations of synapses, with extensive studies conducted in hippocampal CA3-CA1 and CA1-subicular synapses^7,16,17^, along with parabrachial inputs upon the bed nuclei of the stria terminalis^16^, synapses between cone photoreceptors and horizontal cells of the retina^18^, and inputs from the nociceptive pontine parabrachial nucleus onto the central amygdala^19^. The Gβγ-SNARE pathway has also been shown to be of importance outside of the presynaptic terminal, regulating insulin release from beta cells of the islets of Langerhans^20^, as well as large dense core granule release from chromaffin cells^21^. The consequences of chronic, global disruption of the Gβγ-SNARE interaction have been studied in the mouse as a model system. Mutagenesis of SNAP25 to a mutant SNAP25Δ3 that interacts less well with Gβγ leads to physiological consequences, including deficits in stress processing and depressive phenotypes^16,22^.

The binding site for Gβγ on SNAP25 involves two clusters of nine residues on the external surface of the formed SNARE complex, with a well-studied site at the C-terminus^15,23^ that overlaps with, but is distinct from, the primary interface for synaptotagmin 1 binding upon SNAP25 ^24–26^. Truncation of the C-terminal binding site, either via mutagenesis or proteolysis of SNAP25 through botulinum toxin A, reduces the ability of Gβγ to bind SNAP25^13,23,27^. A second site exists adjacent to the N-terminus of the SN1 and SN2 SNARE-forming helices, and the physiological consequences of disruption of this site are unknown^15^. While the regions for the binding sites of Gβγ on SNAP25 have been characterized, much less is known about whether other SNARE proteins also participate in the Gβγ-SNARE pathway. It is known that Gβγ binds to monomeric synaptobrevin and syntaxin1A, the other components of the canonical ternary SNARE complex, in addition to SNAP25. However, given that proteolysis of the C-terminus of SNAP25 abolishes the functional effects of the Gβγ-SNARE pathway in *ex vivo* preparations, what is the contribution of the Qa-SNARE to the pathway? Moreover, numerous genes encoding for Qa-SNAREs and Qb,c-SNAREs exist in the human genome, with 15 syntaxin isoforms^28^, as well as SNAP23^29^, SNAP29^30^, and SNAP47^31^, which have been shown to participate in secretory vesicle trafficking and exocytosis. Can these SNARE-driven processes be regulated by Gβγ, and what is the functional consequence of such regulation?

In this article, we characterize the biochemical binding and functional parameters of four Qa-SNAREs: syntaxins 1-4 and SNAP25, SNAP23, SNAP29, and SNAP47. We assess if they can bind purified Gβγ, participate in lipid mixing as t-SNAREs with the R-SNARE VAMP2, support enhancement of this process by the C2AB-domain Ca^2+^-sensor synaptotagmin I, and whether or not this enhancement can be inhibited by purified Gβγ. These SNAREs have previously been shown to be implicated in a number of critical physiological processes, including insulin release from pancreatic beta cells^32,33^, lysosomal exocytosis^34^, zymogen granule release from the apical surface of pancreatic acinar cells^35,36^, and the fusion of GLUT4-containing vesicles for glucose uptake in skeletal myocytes or adipocytes^37–40^. These SNAREs may participate in the postsynaptic trafficking of ionotropic glutamate receptors^41–43^, a process critical for the induction and maintenance of long-term potentiation in neurons. If biochemical studies could demonstrate the feasibility of Gβγ-dependent modulation of these SNARE proteins, it could provide a mechanistic basis to investigate the role of the Gβγ-SNARE pathway for these processes *in vivo*.

## Results

To conduct biochemical studies, we first sought to express four exocytotic syntaxins in a recombinant manner, as well as four Qb,c-SNARE binding partners. The coding sequences for full-length SNAP23, SNAP29, Stx2 Stx3, and Stx4 were cloned into pGEX6p-1 and expressed in recombinant form in *E.coli*. A truncated fragment of SNAP47 with improved solubility^31^ was also cloned and expressed in a similar manner. To determine if these monomeric SNARE proteins could interact with Gβγ subunits, we utilized the Alphascreen protein-protein interaction assay^27^, in which His-tagged recombinant Gβ1γ2 reacts with biotinylated SNARE proteins in a concentration-dependent manner. As a negative control, biotinylated glutathione-S-transferase was exposed to the highest concentration of Gβ_1_γ_2_ tested to verify that no non-specific binding was generated. All biotinylated proteins were biotinylated at primary amine residues. Biotinylated SNAP23 was found to bind Gβ_1_γ_2_ in a concentration-dependent manner with a significantly lower potency than SNAP25 (**Fig. 1A)**. Biotinylated Stx2 and Stx4 also bound Gβ_1_γ_2_ in a concentration-dependent manner with nanomolar potency **(Fig. 1B, 1C),** consistent with prior biochemical binding studies with Stx1A^13^. We conclude that SNAP23, Stx2, and Stx4 can all bind to Gβγ.

**Figure 1.**
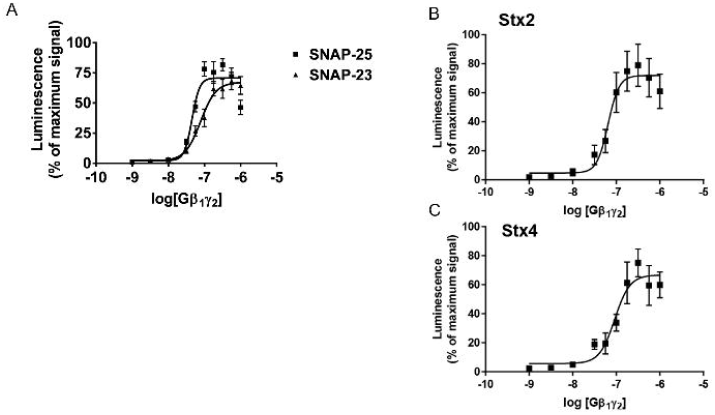
Gβ_1_γ_2_ binds to multiple Qa-SNARE and Qb,c-SNARE isoforms. A. Concentration-response curve for the binding of biotinylated full-length SNAP23 to His-tagged Gβ_1_γ_2_ in the Alphascreen protein-protein interaction assay. The EC50 for the binding of SNAP25 to Gβ_1_γ_2_ is 45.2nM (95% C.I.: 37.2 to 54.9 nM), while the EC50 for the binding of SNAP23 to Gβ_1_γ_2_ is 78.2nM (95% C.I.: 58.6 to 104.8 nM. There were nine total technical replicates per condition prepared from two or more batches of protein. B.: Concentration-response curve for the binding of biotinylated full-length Stx2 to His-tagged Gβ_1_γ_2_ in the Alphascreen protein-protein interaction assay. The EC50 for the binding of Stx2 to Gβ1γ2 is 64.8nM (95% C.I.: 41.5 to 92.1 nM). There were six total technical replicates per condition prepared from two batches of protein. C: Concentration-response curve for the binding of biotinylated full-length Stx4 to His-tagged Gβ1γ2 in the Alphascreen protein-protein interaction assay. The EC50 for the binding of Stx4 to Gβ1γ2 is 92.1nM (95% C.I.: 56.2 to 129.3 nM). There were nine to fifteen total technical replicates per condition prepared from four batches of protein.

The functional consequences of the binding of Gβγ subunits, expressed ubiquitously, to Stx2 Stx3 and, Stx4, as well as SNAP23, SNAP29, and SNAP47, are completely unknown. To determine if Gβγ could inhibit Ca2+-synaptotagmin activity with t-SNAREs formed from these non-neuronal SNAREs, we utilized a reconstituted lipid mixing assay in which liposomes harboring Qa-Qb,c-SNARE complexes were reacted with VAMP2-harboring liposomes **(Fig. 2A),** containing two fluorescently labeled lipids, NBD-phosphatidylethanolamine and rhodamine-phosphatidylethanolamine, which undergo Forster resonance energy transfer (FRET)^44,45^. Upon mixing of the unlabeled t-SNARE harboring liposomes with v-SNARE harboring liposomes, a dilution of the FRET pair occurs, and the normalized NBD fluorescence emitted at 535nm increases. Previously, we have shown that Gβγ inhibits syt I-stimulated SNAP25 and Stx1A-driven lipid mixing in a Ca^2+^-dependent manner^14,16^ **(Figure 2A)**. Furthermore, liposomes harboring t-SNAREs made with syntaxin1 and a mutant SNAP25 that binds less well to Gβγ^27^ are also less sensitive to the inhibitory action of Gβγ^16^ in this assay. We assembled purified preparations of all 16 Qa-Qb,c-SNARE dimers from Stx1A, Stx2, Stx3, and Stx4 paired with SNAP25, SNAP23, SNAP29, and SNAP47 embedded in liposomes. Liposomes containing purified recombinant human syntaxin2 and SNAP25 were reacted with VAMP2-containing fluorescent liposomes with or without 10μM synaptotagmin I 135-421 (syt) in the presence of 100μM Ca^2+^: Since the functional enhancement of syt I and the inhibitory effect of Gβγ upon liposomes harboring syntaxin1-SNAP25 dimers have been characterized by our group in two previous articles^14,16^, those studies were not repeated here. We opted to use purified bovine Gβ_1_γ_1_ for this experiment, as this well-studied heterodimer can be prepared in large quantities in the absence of any detergents which may perturb the lipid mixing assay, or affinity tags, which may alter binding. In the presence of both concentrations of Ca^2+^ and syt, a large increase in NBD-phosphatidylethanolamine fluorescence was observed for liposomes harboring syntaxin2-SNAP25 complexes **(Fig.2B, 2E))**. In the absence of Ca^2+^, a clamping effect of apo-syt was observed relative to conditions lacking syt. 6μM Gβ_1_γ_1_ significantly inhibited syt and Ca^2+^-dependent lipid mixing at 100μM Ca^2+^. Together, this data suggests that syt I can drive lipid mixing using a ternary SNARE consisting of syntaxin2, SNAP25 and VAMP2, and Gβγ can inhibit the stimulatory effect of syt I on this preparation. Next, liposomes harboring syntaxin3 and SNAP25 in complex were tested in an identical fashion. The maximal extent of lipid mixing in conditions with and without Ca^2+^-syt was notably lower for the syntaxin3-SNAP25 pair than syntaxin2-SNAP25 or syntaxin1-SNAP25 (Fig. 2C, 2F). Ca^2+^-syt was able to significantly enhance the maximal extent of lipid mixing with syntaxin3-SNAP25, and apo-syt was able to produce a significant clamping effect relative to conditions lacking syt. Gβγ was able to significantly inhibit the stimulatory effect of Ca^2+^-syt in this preparation. Next, a population of liposomes harboring syntaxin4-SNAP25 complexes was obtained. Liposomes harboring syntaxin4-SNAP25 were able to participate in lipid mixing with VAMP2 (Fig. 2D, 2G), and a stimulatory effect was observed when 100μM Ca^2+^ and syt were present in tandem, but application of 6μM Gβγ did not significantly inhibit the maximal extent of lipid mixing relative to conditions lacking Gβγ. Furthermore, no significant clamping effect of apo-syt relative to conditions lacking syt could be observed. Together, this data suggests that the four Qa-SNAREs tested can all form functional SNARE complexes with SNAP25 that participate in Ca^2+^-syt -dependent lipid mixing, but an isoform-specific role was observed in the regulation of SNAP25-containing SNARE complexes by Gβγ, as Stx4 was insensitive to Gβγ.

**Figure 2:**
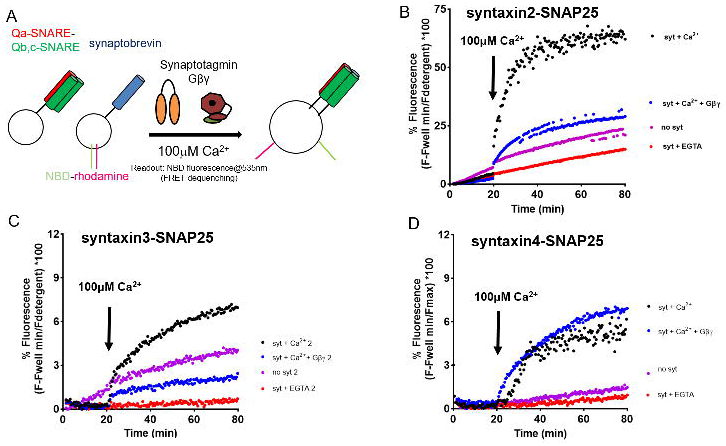
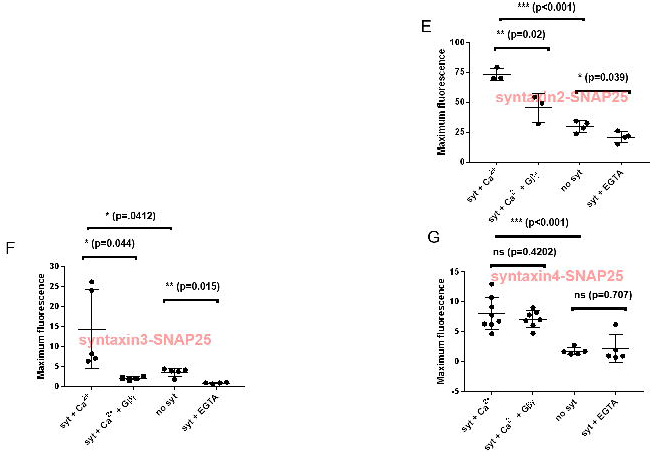
Differential Gβγ sensitivity is observed in lipid mixing with SNAP25 and a panel of syntaxins. A: Diagram showing assay principle. Synaptobrevin-bearing liposomes (containing ~50 copies/vesicle) containing the FRET pair NBD and rhodamine, covalently attached to PE, fuse with unlabeled liposomes containing t-SNARE complexes (containing ~ 130 copies/vesicle) consisting of the tested Qb,c-SNARE and syntaxin isoform. The increased surface area of the fused liposome reduces the quenching of NBD fluorescence by rhodamine and NBD fluorescence increases as a result. B-D: Traces of lipid mixing experiments. At *t* = 20 min, 100 uM CaCl_2_ was added. Fluorescence is normalized by subtracting the well fluorescence and dividing by the maximum fluorescence recorded after addition of detergent. E-G: Graph of maximum fluorescence values for each condition. The specific SNARE protein isoforms described within each graph are denoted in transparent red. E: Bar graph of results obtained from lipid mixing experiments with Syntaxin2-SNAP25 liposomes F: Bar graph of results obtained from lipid mixing experiments with in Syntaxin3-SNAP25 liposomes. G: Bar graph of results obtained from lipid mixing experiments with in Syntaxin4-SNAP25 liposomes. There were three to eight technical replicates per condition prepared from two or more batches of protein. Center bars represent the mean, with small bars representing the S.D of each group.

SNAP23 is the second most widely studied Qb,c-SNARE and is known to participate in many types of exocytosis, some of which are regulated by Gi/o-coupled GPCRs, potentially via the Gβγ-SNARE pathway, but little is known of SNAP23’s ability to participate in lipid mixing^46,47^. Much like in Fig. 2, we conducted an array of lipid mixing studies in a defined liposomal system with SNAP23, along with the four exocytic syntaxins and synaptobrevin. Preparations of liposomes containing SNAP23 in complex with Stx1A, Stx2, Stx3, and Stx4 were obtained. The liposomes containing the Stx1A-SNAP23 complex participated in lipid mixing with synaptobrevin-harboring liposomes (Fig. 3A, 3E) in the absence of syt, and a robust stimulatory effect upon lipid was observed with Ca^2+^-syt, but we were unable to detect a significant clamping effect of apo-syt. Conditions containing 6μM Gβγ along with Ca^2+^-syt showed a near-complete abolishment of the latter’s stimulatory effect. Liposomal populations harboring Stx2-SNAP23 also underwent basal lipid mixing and a large stimulatory effect of Ca^2+^-syt (Fig. 3B, 3F), but distinct from Stx1A-SNAP23, a significant clamping effect of apo-syt upon Stx2-SNAP23 was detected. In conditions with Gβγ, a significant inhibition of the effect of Ca^2+^-syt was again observed. The liposomal populations harboring Stx3-SNAP23 complexes showed a general trend of low activity in all conditions, unlike all previous populations tested(Fig. 3C, 3G). However, we did observe a significant stimulatory effect of Ca^2+^-syt relative to apo-syt or the absence of syt. The presence of 6μM Gβγ in addition to Ca^2+^-syt reversed this effect. A clamping effect was of borderline significance was also observed. Prior studies highlighted a ternary SNARE complex consisting of Stx4, SNAP23, and synaptobrevin as being the active species for GLUT4 vesicle translocation^33,37–40^: we characterized this pair in lipid mixing much like we did for Stx1A, Stx2, and Stx3(Fig. 3D, 3H). Virtually no change in NBD fluorescence whatsoever was detected in any conditions. Since both basal lipid mixing and Ca^2+^-syt lipid mixing with Stx4-SNAP23 were indistinguishable from baseline, we did not test the effect of 6μM Gβγ. Since syt I is not the active species in GLUT4 insertion, but rather Doc2b^48^, we expressed and purified the C2AB domain of human Doc2b^49^ and tested whether it could stimulate Stx4-SNAP23 lipid mixing with synaptobrevin. Ca^2+^-Doc2b produced a negative deflection in NBD fluorescence, but apo-Doc2b was identical to apo-syt and basal conditions lacking any Ca^2+^ sensor. From the totality of these SNAP23 studies, we conclude that SNAP23 can participate in lipid mixing with three of the four exocytic syntaxins tested along with synaptobrevin, that Ca^2+^-syt can exert a stimulatory effect upon this process, and that Gβγ can inhibit this process, much like SNAP25.

**Figure 3:**
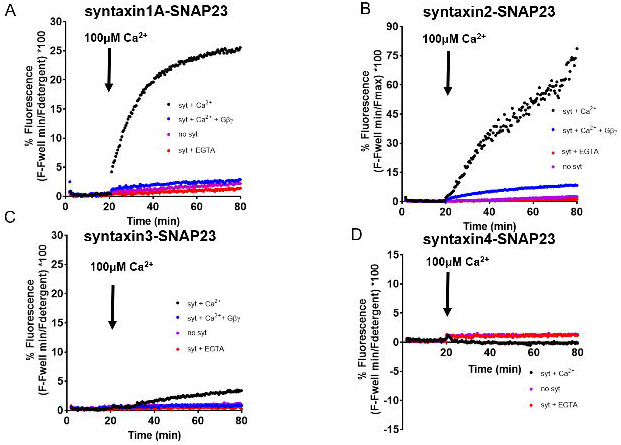
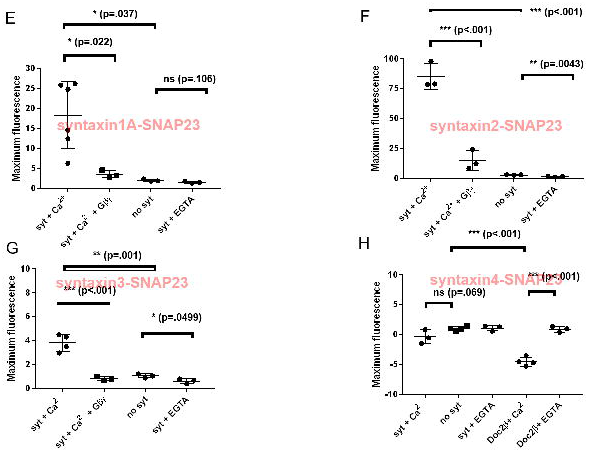
SNAP23-driven lipid mixing can be regulated by Gβγ. A-D: Traces of lipid mixing experiments for four different combinations of SNAP23-harboring liposomes as in Fig. 2. At *t* = 20 min, 100 uM CaCl_2_ was added. Fluorescence is normalized by subtracting the well fluorescence and dividing by the maximum fluorescence recorded after addition of detergent. EG: Graph of maximum fluorescence values for each condition. The specific SNARE protein isoforms described within each graph are denoted in transparent red. E: Bar graph of results obtained from lipid mixing experiments with Syntaxin1-SNAP23 liposomes F: Bar graph of results obtained from lipid mixing experiments with in Syntaxin2-SNAP23 liposomes. G: Bar graph of results obtained from lipid mixing experiments with in Syntaxin3-SNAP23 liposomes. H: Bar graph of results obtained from lipid mixing experiments with in Syntaxin4-SNAP23 liposomes. There were three to eight technical replicates per condition prepared from two or more batches of protein. Center bars represent the mean, with small bars representing the S.D of each group.

SNAP29 is a ubiquitously expressed Qb,c-SNARE isoform required in mammals for normal postnatal development, potentially through its role in intracellular processes such as autophagy^50^. Biochemical functional characterization of SNAP29 has been investigated with VAMP8 and Stx17 ^51^, along with synaptobrevin and Stx1A^46^. However, nothing is known about whether it can participate in the Gβγ-SNARE pathway to inhibit exocytosis. We investigated whether SNAP29 could functionally engage in lipid mixing with our panel of exocytic syntaxins and synaptobrevin, along with syt I. Liposomes containing the previously characterized Stx1A-SNAP29 complex behaved identical to previous studies by other groups^46^, with basal lipid mixing occurring between these liposomes and synaptobrevin-harboring liposomes and a significant enhancement of the rate and maximal effect by Ca^2+^-syt(Fig, 4A,4E). A significant clamping effect was observed upon Stx1A-SNAP29-driven lipid mixing from apo-syt. For the first time, we were able to show a significant inhibitory effect of Gβγ on SNAP29 function in Ca^2+^-syt–dependent lipid mixing. The population of liposomes containing Stx2-SNAP29 behaved similarly to Stx1A-SNAP29 in parameters tested (Fig. 4B, 4F), including the inhibitory effect of Gβγ, continuing a trend observed with SNAP25 and SNAP23. However, these results were not conserved for syntaxin3-SNAP29 complexes embedded in liposomes. While basal lipid mixing was detectable, no stimulatory effect of Ca^2+^-syt could be detected, nor could any inhibitory clamping effect of apo-syt be detected(Fig. 4C, 4G). Since the mechanism of Gβγ is direct competition with syt, the effect of Gβγ upon lipid mixing was not tested for syntaxin3-SNAP29. The fourth SNAP29-containing preparation, syntaxin4-SNAP29, showed a trend with syntaxin4-SNAP25, where basal lipid mixing was observed (Fig. 4D, 4H), and a trend towards a moderate stimulatory effect of Ca^2+^-syt was detected, but significance was not achieved relative to conditions lacking syt, or conditions containing Gβγ. None of the conditions tested for Stx4-SNAP29 were significantly different from any other. From this, we infer that Ca^2+^-syt is less stimulatory for Stx4-SNAP29, and Stx4 as a Qa-SNARE shows no ability to be functionally inhibited by Gβγ.

**Figure 4:**
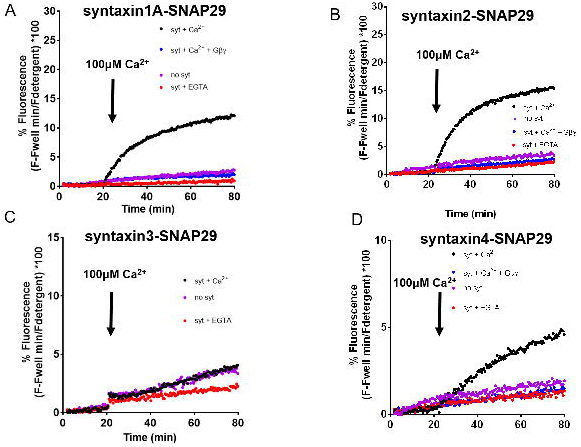
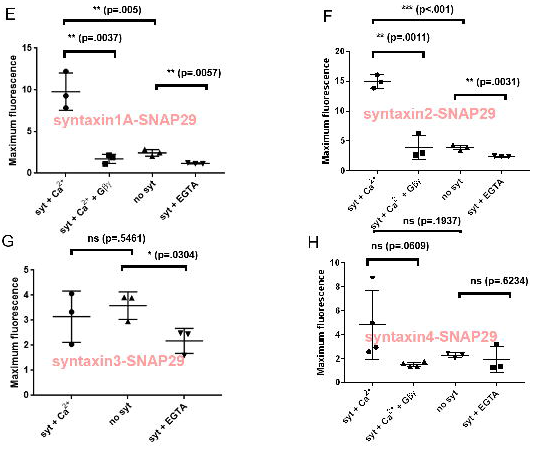
SNAP29-driven lipid mixing can be regulated by Gβγ. A-D: Traces of lipid mixing experiments for four different combinations of SNAP29-harboring liposomes as in Fig. 2. At *t* = 20 min, 100 uM CaCl2 was added. Fluorescence is normalized by subtracting the well fluorescence and dividing by the maximum fluorescence recorded after addition of detergent. EG: Graph of maximum fluorescence values for each condition. The specific SNARE protein isoforms described within each graph are denoted in transparent red. E: Bar graph of results obtained from lipid mixing experiments with Syntaxin1-SNAP29 liposomes F: Bar graph of results obtained from lipid mixing experiments with in Syntaxin2-SNAP29 liposomes. G: Bar graph of results obtained from lipid mixing experiments with in Syntaxin3-SNAP29 liposomes. H: Bar graph of results obtained from lipid mixing experiments with in Syntaxin4-SNAP29 liposomes. There were three to four technical replicates per condition prepared from two or more batches of protein. Center bars represent the mean, with small bars representing the S.D of each group.

SNAP47 is a Qb,c-SNARE with a large N-terminal domain. Biochemical functional characterization of SNAP47 has been limited to a single article^31^. A fragment containing the SNARE domains of SNAP47, termed SNAP47_115-412_, was obtained. We obtained liposomes harboring SNAP47_115-412_ in complex with Stx1A, Stx2, Stx3, and Stx4. These liposomes were tested in a lipid mixing paradigm identical to Figure 2-4. However, no stimulatory effect of syt nor inhibitory effect of apo-syt could be detected in all SNAP47 lipid-mixing experiments, with low basal lipid mixing overall(Fig. 5A-5H). From this, we conclude that the combination of SNAP47, Stx1A, 2, 3, or 4, and synaptobrevin is of low activity in lipid mixing.

**Figure 5:**
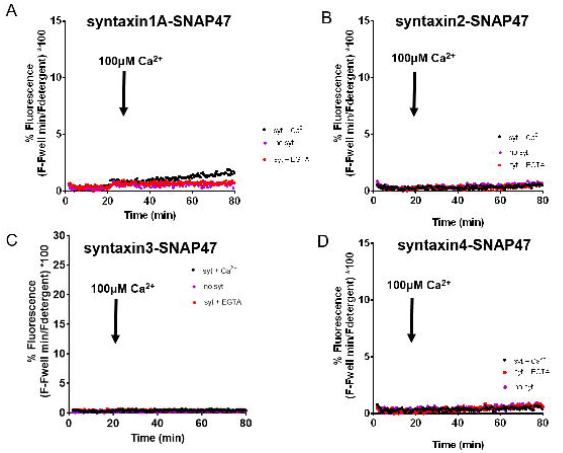
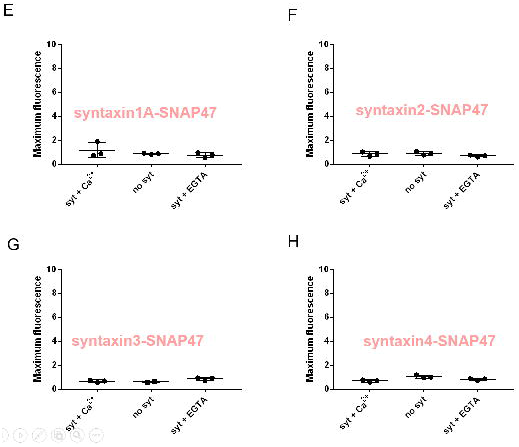
SNAP47_115-412_ is minimally active in lipid mixing with Stx1A, Stx2, Stx3, Stx4, and synaptobrevin. A-D: Traces of lipid mixing experiments for four different combinations of SNAP47-harboring liposomes as in Fig. 2. At *t* = 20 min, 100 uM CaCl_2_ was added. Fluorescence is normalized by subtracting the well fluorescence and dividing by the maximum fluorescence recorded after addition of detergent. E-G: Graph of maximum fluorescence values for each condition. The specific SNARE protein isoforms described within each graph are denoted in transparent red. E: Bar graph of results obtained from lipid mixing experiments with Syntaxin1-SNAP47 liposomes F: Bar graph of results obtained from lipid mixing experiments with in Syntaxin2-SNAP47 liposomes. G: Bar graph of results obtained from lipid mixing experiments with in Syntaxin3-SNAP47 liposomes. H: Bar graph of results obtained from lipid mixing experiments with in Syntaxin4-SNAP47 liposomes. There were three technical replicates per condition prepared from two or more batches of protein. Center bars represent the mean, with small bars representing the S.D of each group.

To identify potential regions of interest for Gβγ binding on our array of SNARE proteins, we performed ResPep peptide mapping studies^15^. In this technique, 15-mer polypeptides corresponding to the primary sequence of a SNARE protein are chemically synthesized and spotted onto a PVDF membrane. Each subsequent peptide advances by 3 C-terminal residues to the previous peptide. Peptides corresponding to syntaxin 6 and 11, the latter of which is involved in platelet exocytosis^52^, but lacks a TM domain, making it challenging to use in recombinant form for lipid mixing studies, were also characterized using this method. The derivatized membrane was then exposed to recombinant His-tagged Gβ1γ2 in a Far-Western blotting procedure(**Fig. 6A)**, where unbound Gβγ was removed by washing and quantitatively detected via chemiluminescence after labeling with primary anti-Gβ antibodies and HRP-conjugated secondary antibodies. As a positive control, a known strongly Gβγ-binding peptide corresponding to the C-terminus of GRK2^53^ was spotted onto the membrane, and a region was left underivatized to act as a negative control. In this procedure, peptides which bind Gβγ strongly generate luminescent signals. First, peptides corresponding to SNAP23 did not show nearly as much interaction with Gβγ relative to SNAP25^15^, which previously highlighted multiple regions of robust binding (**Fig. 6G)**. SNAP23 130-150, at the N-terminus of the SN2 helix, displayed the most robust interaction of any region on SNAP23, with weaker interactions occurring at a region corresponding to residues 18-38. Peptides corresponding to the C-terminal region of SNAP23 did not generate significant Gβγ binding, unlike SNAP25. Next, peptides corresponding to Stx2 113-130, located within the helical Habc region of the protein, strongly interacted with Gβγ(**Fig. 6C)**, as did a second region corresponding to Stx2 144-160. Minimal interaction was observed in the SNARE domain, in contrast to previous studies, in which truncated syntaxin1 SNARE domains bound Gβγ but isolated Habc domains did not^13^, potentially as a result of misfolding of the truncated Habc fragment. The homologous region to Stx2 144-160 on Stx3, 145-161, showed approximately 5-fold weaker binding, while Stx4 152-168 also showed Gβγ binding, with a rank order of 2 > 3 = 4 for this region. While syntaxin6 shows much less homology to syntaxin2, Gβγ binding was predicted for residues 82-105 and 136-156. The platelet syntaxin, syntaxin11, whose post-translational modifications make it challenging to characterize accurately in in vitro binding assays^47^, displayed the most interactivity towards Gβγ of any syntaxin characterized in this experiment, with 3 critical interaction sites at residues 61-90, 115-147, and 208-234.. We conclude that the Gβγ-interacting region on SNAP23 may be less extensive than SNAP25, and that a region in the Habc domain of syntaxins may be important for the interaction with Gβγ.

**Fig. 6:**
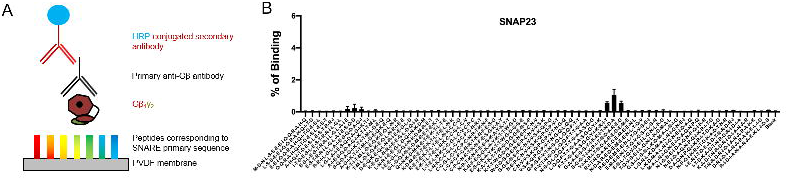
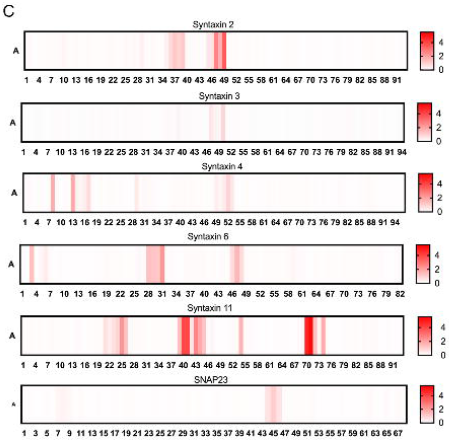
Mapping the binding sites of Qa-SNAREs and Qb,c-SNAREs with ResPep. A. Diagram of Far-Western assay principle. B. Bar graph showing complete array of 15-mer peptides corresponding to the primary sequence of SNAP23 on the X-axis and the percent Gβ_1_γ_2_ binding relative to the most intense spot on the membrane. C. Heat map of the binding site of each Qa-SNARE isoform and Qb,c-SNARE isoform analyzed in this fashion, with areas of high binding shown in red. Error bars represent mean + S.E.M.

## Discussion

Here, we have expanded upon our existing understanding of the Gβγ-SNARE pathway, generalizing previous findings found with discrete SNARE protein isoforms expressed in the nervous system to the more broad family of Qa- and Qb,c-SNAREs. All of the SNARE proteins tested in the Alphascreen assay were able to bind Gβγ. and we speculate that this binding may be a general function shared across all members of the family. While we were unable to demonstrate functional inhibition of Stx4-SNAP25 or Stx4-SNAP29 activity by Gβγ in this study, we cannot rule out the possibility that such activity exists in secretory cells that express both Stx4 and SNAP25 (or the ubiquitously expressed SNAP29), such as pancreatic beta cells or numerous populations of neurons.

### Differential basal lipid mixing and Ca^2+^-syt function between between Qb,c-SNARE isoforms

We were unable to detect significant lipid mixing between liposomes harboring any combination of Stx1A, Stx2, Stx3, or Stx4 along with SNAP47_115-412_ and liposomes harboring synaptobrevin. A single research group was able to detect a low level of basal lipid mixing between Stx1A, SNAP47_115-412_ and synaptobrevin^31^. While slight differences in the way the assay is performed may affect functional results, these studies share a conclusion that SNAP47 shows less activity in this type of assay than SNAP25 or SNAP23. The combination of Stx3 and Stx4 with SNAP29 also showed less activity with synaptobrevin than Stx1 or Stx2 with SNAP29. A tandem role for Stx3 and Stx4 with SNAP29 in secretory autophagy has previously been highlighted^54^. These functions may require a different R-SNARE such as VAMP8. For pairs that showed low activity in this study, more robust activity may be observed with other R-SNARE isoforms, which were beyond the scope of our approach.

### *SNAP23 and SNAP29 bind to and are functionally inhibited by* Gβγ

We have shown here that SNAP23 binds Gβγ, albeit it somewhat less well than SNAP25 does. We are in the process of studying SNAP29 binding to Gβγ. This is consistent with binding studies showing incomplete conservation of known SNAP25 Gβγ-binding residues on SNAP23. Seven known Gβγ-binding residues are fully conserved-E62, K102, R135, R136, R142, R198, and K201^15^. However, the extreme C-terminus is not conserved, with a SNAP23 containing a bulky, negatively charged Asp residue instead of Gly for SNAP25. Given that the G204* mutation reduces SNAP25’s ability to bind Gβγ by twofold^27^, reduces the ability of Gβγ to inhibit SNAP25-driven lipid mixing^16^, and disrupts the ability of G_i/o_-coupled GPCRs to inhibit exocytosis^16^, we would anticipate that the presence of this Asp for SNAP23 would be disruptive to the local binding environment. A number of SNAP23 and SNAP29-harboring pairs showed strong inhibition by Gβγ in this study. More broadly, any cellular secretory process that involves SNAP23 or SNAP29 is potentially a candidate for regulation by free cellular Gβγ or any Gi/o-coupled GPCR that works via the Gβγ-SNARE pathway.

### *Variable interaction with* Gβγ *as a trait of Qa-SNARE isoforms*

Here, we have provided evidence that interaction with Gβγ is a feature shared widely between members of the syntaxin family. Syntaxin2 and 4 were shown to bind Gβγ as full-length monomers, and peptides derived from syntaxin 3, 6, and 11 were also shown to bind Gβγ. Functional regulation of syntaxin2 activity was established **(Fig. 2-3)**, although we were unable to demonstrate such regulation for syntaxin4 **(Fig. 4)**. Despite identical preparations, extent of SNAP25 binding, and concentrations utilized, t-SNAREs containing syntaxin4 behaved much differently than Stx1A, Stx2, or Stx3 in lipid mixing studies **(Fig. 3-4)**, with a substantially reduced and slower maximal extent of lipid mixing for syntaxin4 than syntaxin2, both in the presence and absence of syt I. The fact that we could not establish a role for Gβγ in inhibiting the syntaxin4-dependent mechanism of lipid mixing suggests that this process may proceed through a different mechanism than syntaxin2-dependent or Stx1A-dependent lipid mixing. We were unable to show a clamping effect by apo-syt for Stx4-SNAP25 liposomes, and Stx4-SNAP29 liposomes trended towards significance with regard to a stimulatory effect of Ca^2+^-syt, potentially suggesting that the two syntaxins have different syt-binding properties. In the X-ray crystallographic structure of syt I binding to the ternary SNARE complex^24,25^, three residues on Stx1A are seen in the primary interface: D231, E234, and E238. E238, in particular, is not conserved between the three syntaxins for which we have characterized lipid binding studies: the corresponding residue is a Met residue for Stx2, and a Leu residue for Stx4. The substitution of a charged, highly polar glutamate residue for a very hydrophobic leucine residue may provide insights for future investigation in differential function between the two syntaxin isoforms. Syntaxin11, investigated here as a series of peptides **(Fig. 5)** bound more Gβγ than any of the four other isoforms tested in this assay. Syntaxin11 has been previously shown to be the most important Qa-SNARE involved in α–granule and dense granule exocytosis^52^, as demonstrated by *ex vivo* secretion studies in platelets from human familial hemophagocytic lymphohistiocytosis type 4 (FHL4) patients lacking syntaxin11. Unfortunately, syntaxin11 lacks a transmembrane domain and is dependent on posttranslational acylation for membrane anchoring, rendering it challenging to produce in the native state via recombinant methods^47^. We observed the least amount of Gβγ binding for syntaxin3 of any Qa-SNARE isoform tested, which only showed a small amount of binding in the Habc region. Syntaxin3 has been implicated in the exocytosis of von Willebrand factor within Weibel-Palade bodies from endothelial cells in humans^55^, and variant microvillus inclusion disease(MVID) patients have been shown to have null mutations in both copies of syntaxin3^55,56^. However, robust Gβγ function was observed in lipid mixing for Stx3-SNAP25 and Stx3-SNAP23, implying that this small interaction site is sufficient.

## Materials and Methods

### Protein Purification

Plasmids for GST tagged SNAP23, SNAP29, SNAP47_115-412_, Syntaxin 2, Syntaxin3, Syntaxin 4, and Synaptotagmin 1A, as well as HIS tagged Syntaxin 1 and SNAP-25 were transformed with BL21-DE3 bacteria. Bacteria were grown at 37 C overnight in a starter culture overnight with a 1:1000 dilution of ampicillin. SNAP-25 and syntaxin 1 used 1:1000 dilution of kanamycin. The starter culture was then transferred to 2 1 L flasks with the same dilution of either ampicillin or kanamycin and grown until an A_600_ of 0.8 was reached. Then, the bacteria was either induced with 0.4 mM isopropyl β-D-thiogalactoside (IPTG) for 4 hours at 37 C or overnight at 25 C for the SNAP-25, Syntaxin 1, and Synaptotagmin 2. The bacteria was then pelleted and resuspended in resuspension buffer. The resuspension buffer always consisted of protease inhibitors (phenylmethylsulfonyl fluoride, leupeptin, aprotinin, pepstatin). In addition, for SNAP-23, the buffer consists of 50 mM HEPES pH 8.0, 140 mM KCl, 5 mM 2-mercaptoethanol, and 1 mM EDTA. The syntaxins had the same buffer but with 25 mM HEPES pH 8.0 and 400 mM KCl. For SNAP-25, the buffer consisted of the protease inhibitors, 50 mM Tris pH 7.8, 1% Triton X-100, 5 mM beta mercaptoethanol, and finally 500 mM sodium chloride. Finally the synaptotagmin has a resuspension buffer consisting of the protease inhibitors and 40 mM Tris pH 8.2, 200 mM NaCl, 1% Triton X-100, and 5 mM 2-mercaptoethanol. After sonication, the bacteria was put in the ultracentrifuge for 30 minutes at 22,000 RPM (Type 70 Ti). For the syntaxins, Triton X-100 was added to achieve a final concentration of 20%. The syntaxins were then put on the rotor (Ti-70) at 4 C for 3-4 hrs. They were then ultracentrifuged at 36,000 RPM for 1 hour. In the meantime, 2.5 mL of Glutathione Agarose Beads or 2 mL of TALON Metal Affinity Resin for HIS tagged proteins were washed with resuspension buffer (2 times with 5 minutes on 4 C rotor and centrifuged at 4000 rpm for 6 minutes). The lysate from the ultracentrifuge was then added to the beads and incubated at 4 C on the rotor overnight. Afterwards, the beads were washed (same washing procedure as before) three times with wash buffer which, for SNAP 23 and 25, consisted of 25 mM HEPES pH 8.0, 150 mM KCl, 5 mM 2-mercaptoethanol, 1% Triton X-100, and 1 mM EDTA. For the syntaxins, the buffer consisted of 25 mM HEPES pH 8.0, 400 mM KCl, 5 mM 2-mercaptoethanol, 1 mM EDTA, and 1% Triton X-100. 2 mM magnesium chloride was added to the Synaptotagmin 2 during the washes to remove any nucleic acids. The beads were then washed (same washing procedure as before) with elution buffer (same as wash buffer but with 10% glycerol and 20% n-Octylglucoside instead of Triton X-100) four times before being resuspended in elution buffer. A 1:100 dilution of 3C-Protease was then added and incubated on a 4 C rotor overnight for GST tagged proteins. HIS tagged proteins were cleaved by adding aaa and incubating at 37 C for two hours. Finally, the beads were spun down and the supernatant containing the protein was extracted. SDS-Page was performed on the proteins to confirm their purity. The bicinchoninic acid assay was also used (following manufacturer’s instructions) to determine protein concentration.

### Liposome Preparation

Lipids consisting of 55% POPC (1-palmitoyl-2-Oleoyl-sn-glycero-3-phosphocholine), 15% DOPS (1,2-dioleoyl-sn-glycero-3-phospho-L-serine (sodium salt) and 30% POPE (1-palmitoyl-2-oleoyl-sn-glycero-3-phosphoethanolamine) were dried down using nitrogen then a vacuum desiccator. The next morning, they were resuspended in 500 uL elution buffer (25 mM HEPES-KOH pH 7.8, 400 mM KCl, 10 % glycerol, 5 mM 2-mercaptoethanol, 1 % n-Octylglucoside). The appropriate isoform of SNAP and syntaxin proteins were added to the solution to reach the desired concentration. After vortexing gently for 15 minutes, 2 volumes of elution buffer was slowly added. The tubes were then vortexed gently for another 15 minutes. The solution was transferred to dialysis tubing which was pre soaked with water, then elution buffer. Two 4 L beakers with dialysis buffer (25 mM HEPES pH 7.8, 100 mM KCl, 10 % glycerol, 1 mM DTT) was used. After dialyzing in the first beaker for 6 hours, the tubing was then transferred to the other beaker to dialyze overnight. The liposomes were then floated by using a density gradient consisting of 80%, 30%, and 0% Accudenz. The gradient was assembled in centrifuged tubes for the SW-55 rotor. The tubes were then centrifuged at 55,000 rpm for 2 hours at 4 C. The liposomes between the 0 and 30 percent gradient were then harvested and stored at −80 C.

### Lipid Mixing Assay

Liposomes were thawed and the synaptotagmin was added. In addition, Gβγ was added if required. EGTA was then added and buffer consisting of 25 mM HEPES pH 7.4, 100 mM KCl, and 1 mM DTT was added to reach a final volume of 75 uL. Once in the wells, the v-SNAREs were added. The plate was then placed in the plate reader at 37 C. The experiment was left to run for 20 minutes to let all calcium independent fusion occur. Ca^2+^ was added after 20 minutes and then left to run for one hours. Finally, n-octylglucoside was added and left to sit for 15 minutes.

### Peptide array synthesis

Peptide array synthesis was performed using the Respep SL (Intavis AG) according to standard SPOT synthesis protocols^57^ according to previously published methods^15^. The Respep SL system managed complex timing, mixing, additions, and washing of the membrane. Peptides were 15 residues in length and derived from primary sequences of human SNAP23, Stx2, Stx3, Stx4, Stx6, and Stx11 on the sequence available from the UniprotKB/Swiss Protein Database. Peptides were synthesized coupled to the membranes and membranes were the processed with a side chain deprotection step. Membranes in a solution of 95% trifluoroacetic acid, 3% tri-isopropylsilane for 1 hour under agitation. After removal of trifluoroacetic acid, the membrane was then washed: first with dichloromethane, then dimethylformamide, followed by ethanol, and dried completely.

### Peptide membrane Far-Western

Dried membranes were re-hydrated over two washes for 5 minutes in water. The membranes were blocked from non-specific interactions for 1 hour in a buffer of tris-buffered saline (TBS) with 5% milk and 0.1% Tween-20 (Sigma-Aldrich) under gentle agitation, followed by washing 5 times for 5 minutes in TBS with 0.1% Tween-20 on a shaker at RT. For the binding step, membranes were incubated overnight at 4°C with Gβ_1_γ_2_ at a final concentration of 0.44 μM in a binding buffer of 20 mM HEPES, pH 7.5, and 5% glycerol, and washed three times at room temperature on a shaker for 5 minutes in TBS with 5% milk and 0.1% Tween-20. In the primary antibody step, membranes were then exposed to anti Gβ (T-20, SC-378; Santa Cruz Biotechnology, Inc.) at 1:5000 dilution in TBS with 0.2% Tween-20 with mixing for 1 hour before being washed three times for 5 min in TBS with 0.1% Tween-20. HRP-conjugated anti-rabbit secondary antibody was then diluted into TBS to 1:10,000 dilution with 5% milk and 0.2% Tween-20 followed by gentle agitation on a shaker with membranes for 1 hour. Membranes were then washed twice for 5-minutes in TBS with 0.1% Tween-20, followed by two 10-minute washes in TBS. Membranes were imaged via chemiluminescence.

### Data Analysis

The minimum fluorescence during the first phase of the experiment was subtracted from the average fluorescence post calcium addition. Finally, these values were divided by the maximum value from the post-detergent addition stage.

## Acknowledgements

We would like to acknowledge the laboratory of Josep Rizo at the University of Texas-Southwestern for the SNAP25 and synaptotagmin 135-421 constructs and the laboratory of Edwin R. Chapman at the University of Wisconsin-Madison for assistance in conducting assays. We would like to thank VectorBuilder for assistance in developing the SNAP29, SNAP47_115-412_, and Stx3 GST-fusion bacterial expression constructs. We would like to thank Joel H. Everett for assistance in performing protein expression and Alphascreen assays.

